# QTL analysis of vegetative phase change in natural accessions of *Arabidopsis thaliana*

**DOI:** 10.1101/2021.10.27.465806

**Authors:** Erin Doody, Yuqi Zha, Jia He, Scott Poethig

## Abstract

Shoot development in plants is divided into two phases, a vegetative phase and a reproductive phase. Vegetative growth also has two distinct juvenile and adult phases, the transition between which is termed *vegetative phase change*. To understand how this developmental transition is regulated in natural populations of plants, we grew a group of 70 accessions of *Arabidopsis thaliana* and measured the appearance of traits associated with vegetative and reproductive phase change. We found that these transitions were uncorrelated, implying they are regulated by different mechanisms. Furthermore, an analysis of accessions from Central Asia revealed that precocious changes in leaf shape poorly correlated with the timing of abaxial trichome production (an adult trait) and with variation in the level of miR156 (a key regulator of vegetative phase change). This suggests the timing of vegetative phase change is regulated by more than one mechanism. To identify the genes responsible for the precocious vegetative phenotype of these accessions, we used a set of recombinant inbred lines derived from a cross between the standard lab strain, Col-0, and one of these accessions, Shakdara. We identified eight quantitative trait loci involved in the vegetative phase change, some of which regulated different components of leaf development. All of these loci were distinct from those that regulate flowering time. These data provide the foundation for future studies to identify the loci and the regulatory networks responsible for natural variation in the timing of vegetative phase change in *A. thaliana*.

## Introduction

In land plants, shoot growth can be divided into several distinct developmental phases. Following gemination, a plant begins a period of juvenile vegetative growth, a developmental phase that can be very short or last years (Wareing 1959). It then undergoes vegetative phase change and transitions to adult vegetative growth. The transition to the reproductive phase usually occurs after this (although it can also occur during the juvenile phase in some species), and is accompanied by the production of structures involved in sexual reproduction, such as flowers or cones (Levy and Dean 1998; Amasino 2010; Huijser and Schmid 2011; Andrés and Coupland 2012). Finally, to complete its life cycle, the plant undergoes senescence and dies (Leopold 1961; Thomas *et al.* 2003). The duration of these phases is highly variable between and within plant species, and transitions between them are dependent on both development pathways and environmental cues (Conway and Poethig 1993).

Vegetative phase change, or the transition from juvenile to adult vegetative growth, is accompanied by changes in shoot physiology and morphology, including immune and defense responses, stress responses, and metabolic processes (Allsopp 1967; Hackett 1985; Poethig 1990; Boege and Marquis 2005; James and Bell 2001; Stief *et al.* 2014; Cui *et al.* 2014; Nguyen *et al.* 2017; Leichty and Poethig 2019; Lawrence *et al.* 2021). These changes help the plant become better suited to environmental conditions it encounters as it ages. In *Arabidopsis thaliana*, the most obvious morphological markers of vegetative phase are a decrease in leaf base angle, the production of trichomes on the abaxial surface of the leaf blade, and an increase in number and size of serrations of the leaf margin (Telfer et al., 1997).

The master regulators of vegetative phase change are the pair of microRNAs miR156 and miR157 (Wu *et al.* 2009). miR156/7 are highly expressed in leaves produced early in development and at much lower levels in leaves produced during the adult phase (He et al. 2018). This decrease allows for an increase in their targets, the *SQUAMOSA PROMOTER BINDING-LIKE (SPL)* transcription factors, which in turn regulate many different processes associated with shoot and leaf development (Wang and Wang 2015; Xu *et al.* 2016). Regulation of vegetative development by the miR156/SPL module is conserved across all land plants of varying complexity and evolutionary age, from mosses and liverworts to other long-lived perennials such as trees (Mishler 1986; Wang *et al.* 2011; Tsuzuki *et al.* 2019; Zhang *et al.* 2020; Aloni 2021).

While many aspects of the regulation of vegetative phase change by the miR156/7 pathway have been uncovered through a molecular genetic approach, many questions remain unanswered. In particular, the genetic basis for intra-and interspecies variation in the timing of vegetative phase change, as well its relationship to the reproductive phase transition is unknown (Goebel 1900; Wiltshire *et al.* 1991; Hudson *et al.* 2014; Foerster *et al.* 2015). Studies of natural variation in *A. thaliana* have been vital for understanding the mechanism of flowering time (Alonso-Blanco *et al.* 1998; Lempe *et al.* 2005; Shindo *et al.* 2005; Rosas *et al.* 2014), germination (Chiang *et al.* 2009; Martínez-Berdeja *et al.* 2020), senescence (Lyu *et al.* 2018; He *et al.* 2018b) and many other phenomena (Bergelson and Roux 2010; Yuan and Kessler 2019; Rubin *et al.* 2019; Sasaki *et al.* 2019; Nakano *et al.* 2020; González *et al.* 2020). However, the extent to which the timing of vegetative phase change varies within this species, and the genetic basis for this variation remain to be investigated.

To begin to unravel these questions, we grew a group of 70 genome-sequenced natural accessions of *A. thaliana* in uniform conditions and characterized variation in vegetative phase change. This revealed that the timing of vegetative phase change and reproductive phase change were decoupled in these accessions. We then performed a more detailed analysis of a group of accessions with precocious leaf shape that were collected in Central Asia from environments associated with high variation in temperature. We found different phase-specific vegetative traits were not correlated with each other, or with the expression level of miR156 or *SPL* genes, in these accessions. To further explore the genetic basis of vegetative phase change, we conducted quantitative trait loci (QTL) analysis using a set of recombinant inbred lines (RILs) derived from the accessions Col-0 and Sha. This revealed eight major QTLs involved in this transition, and several promising candidate genes within these loci. Our results indicate that genetic regulation of vegetative phase change in natural populations of *A. thaliana* is complex, and is unlikely to be attributable entirely to variation in expression of miR156 or its targets.

## Materials and methods

### Plant material and growth conditions

Stocks of natural accessions were obtained from Arabidopsis Biological Resource Center (ABRC, Ohio State University) and corresponding geographic data was obtained from the 1001 Genomes Database (Alonso-Blanco *et al.* 2016). ABRC stock numbers of these accessions and geographic coordinates of their collection locations are available in **Table S2**. Seeds were sown on Farfard #2 soil (Farfard) treated with fertilizer (Peters Peat Lite 15-16-17), Marathon 1% granular (Marathon) and Scanmask Spray (Biologic) for insect control. Seeds were kept at 4°C for 2 days (non-vernalized) or four weeks (vernalized) and then transferred to a growth chamber, with the transfer day counted as the planting date. Plants were grown at 22°C under a combination of cool white and Gro LITE fluorescent bulbs (Interlectric, F32T8/GL/WS) in either long day (LD) (16hrs. light/8 hrs. dark; 100 μmol m-2 s-1) or short day (SD) (10 hrs light/14 hrs dark; 130 μmol m2 s-1) conditions. Unless indicated otherwise, all experiments in SD and LD conditions were in four-week vernalized conditions. In experiments with numerous flats, seeds of each genotype were dispersed into multiple groups across flats, and flats were rotated in the growth chambers every three days to facilitate even distribution of light throughout the experiment. No statistical difference between the phenotype of Col-0 plants grown in each flat (Student’s *t*-test), allowing for pooling of data between flats.

For QTL mapping, the 13RV Core-pop set derived from Sha and Col-0 accessions was obtained from the Versailles Arabidopsis Stock Center (Simon et al. 2008, http://publiclines.versailles.inrae.fr/rils/index). 10-12 individuals from these 164 lines, plus the Sha and Col-0 parental lines, were grown as described above in both LD and SD conditions and scored for number of leaves lacking abaxial trichomes (both SD and LD), leaf three base angle (both LD and SD), leaf seven serrations (SD only) and days to first open flower (LD only). There was no statistical difference between the phenotype of Col-0 or Sha plants grown in each flat (Student’s *t*-test), allowing for pooling of data between flats. All phenotypic and genotypic data used for QTL mapping are available in **Table S3**.

### Leaf shape and phenotypic measurements

Juvenile leaf number was scored by counting number of leaves lacking trichomes on the abaxial side of the lamina. Flowering time was measured by the number of days to the opening of the first flower. To analyze leaf shape, fully expanded leaves were attached to paper using double-sided tape and scanned. The angle of the leaf base corresponds to the angle between two lines tangent to the base of the lamina intersecting at the petiole and was measured using Fiji (Schindelin *et al.* 2012). We measured leaf three in RILs, and leaf four in natural accessions. Leaf serrations were counted on leaf seven using the leaf shape analysis program Lamina (Bylesjö *et al.* 2008). In all cases, at least 6-12 plants of each genotype were measured.

### Principal component analysis (PCA) of climatic variables

Collection locations (Longitude and Latitude) of the 70 accessions used in this study were obtained from the 1001Genomes database (https://1001genomes.org/accessions.html). Climate data, which included 19 bioclimatic variables and altitude for each of these locations, were extracted from the WorldClim Database with 30 arc-second spatial resolution (http://www.worldclim.org/current) (Fick and Hijmans 2017). PCA was done in R with the prcomp (center and scale = TRUE) function using the data for these 20 climatic variables. Data are available in **Table S2**.

### Measuring miRNA and mRNA abundance by RT-qPCR

Tissue from whole 2-week old seedlings and leaf primordia 2mm in length was collected in biological triplicates and ground in liquid nitrogen. Total RNA was extracted using Trizol (Invitrogen) and treated with RNAse free DNAse (Quiagen) as per the manufacturers’ instructions. 300ng of total RNA was used for reverse transcription using Superscript III First Strand Synthesis System (Invitrogen). RT-PCR for smRNAs was done using primers specific for mature sequences of miR156 in combination with the stem-loop RT primers described in (Varkonyi-Gasic *et al.* 2007) and reverse primer specific to SnoR101 (Xu *et al.* 2015). For mRNA RT-PCR amplification, we used a polyT primer. The resulting cDNA was quantified by qPCR using a three-step amplification protocol and SYBR-Green Master Mix (Bimake) in technical triplicates with primers specific to each mRNA or miRNA using either SnoR101 (miRNA) or *ACT2* (mRNA) as the endogenous controls (**Table S4**). Relative abundance was calculated using the 2^-ΔΔCt^ method (Livak and Schmittgen 2001). The data presented represent the averages of biological triplicates.

### QTL analysis

QTL interval mapping was implemented in the R/qtl package (Broman *et al.* 2003) in R (https://www.r-project.org/) using phenotypic data from Sha x Col-0 RILs (**Table S3**). Logarithm of the odd (LOD) scores for each phenotype were computed via the ‘scanone’ function based on maximum likelihood via the expectation–maximization (EM) algorithm (black lines), Haley-Knott regression (blue lines), and the multiple imputation method (red lines) (Churchill and Doerge 1994; Sen and Churchill 2001). The experiment-wide 0.05 significance LOD score threshold was determined for each trait from 1000 permutations. Analysis of epistasis for each QTL was performed using the multiple imputation method by including the location of the QTL peak as a covariate to the trait of interest. In short, the ‘scanone’ function was used to scan the genome for associations while controlling for the marker location nearest to each peak as an interacting covariate. All methods are described in more detail here: https://rqtl.org/rqtltour.pdf and Broman and Sen 2009.

### SNPeff

Genomic sequences and annotation files of the Tair10 Col-0 assembly (https://www.arabidopsis.org/download/) and the 1001 Genome Project Sha assembly (https://1001genomes.org/data/MPIPZ/MPIPZJiao2020, (Jiao and Schneeberger 2020) were used for analysis.

Col-0 and Sha whole-genome sequences were aligned and visualized using Mauve alignment software (Darling *et al.* 2004). Homologous regions from the Sha and Col-0 genomes that corresponded to the RIL genomic markers closest to boundaries of each QTL were extracted for further analysis. Chromosome two coordinates used were 7650461-13471565 and 7424367-13257809 for Col-0 and Sha, respectively. Chromosome five coordinates used for analysis were 16368385-27316966 and 16482000-27639917 for Col-0 and Sha, respectively. The resulting sequences were aligned using MAFFT (Katoh, 2002) and converted to variant call format (VCF) files (Lindenbaum, 2015) using Col-0 (Tair10) as the reference sequence. The predicted effects of SNPs on gene function within the QTL regions on chromosomes two and five were then predicted with SnpEff (Cingolani *et al.* 2012). Results are shown in **Table S1**.

### Differential gene expression

Transcription data from leaf tissue were obtained from the 1001 Epigenomes database (1001epigenomes.org) for samples Col-0 (Sample 6909, <u>GSM2136441</u>) and Sha (Sample 10015, <u>GSM2135725</u>) (Kawakatsu *et al.* 2016). Normalized data were filtered to select for genes within each QTL region (same as above) and differential expression was determined using the DeSeq2 package (Love *et al.* 2014) in R (https://www.r-project.org/). Results are shown in **Table S1**.

### Statistical Analysis

All statistical tests and analyses were conducted in R version 3.6.1 (https://www.r-project.org/). Linear regression was done with the lm() function. Student’s *t*-test was done using the stat_compare_means (method = “t.test”) function as part of the library(ggpubr) package. ANOVA was done using the aov(model) function and a linear model (lm() function) between variables. Tukey Honest Significant Differences was done to each pair of treatments using the TukeyHSD(conf.level=0.95) function.

## Results

### Natural accessions of Arabidopsis thaliana show variation in the timing of vegetative phase change

To survey the phenotypic diversity in the timing of vegetative phase change in natural accessions, we selected a group of 70 accessions to represent the geographic and ecological diversity of this species and grew them under SD and LD conditions (Fig. 1A). Plants were scored for the timing of vegetative phase change by measuring the angle of the base of the lamina and by counting the number of leaves lacking abaxial trichomes. Because abaxial trichome production is qualitative trait, the node at which the shoot switches from the juvenile (absent) to the adult (present) form of this trait is easy to determine. In contrast, the angle of the leaf base becomes gradually smaller from leaf to leaf, making it difficult to specify a single node at which the juvenile-to-adult transition occurs. We measured the angle of the leaf base of leaf four on the assumption that accessions with a small leaf base angle at this node undergo vegetative phase change earlier than accessions with a larger leaf base angle.

**Figure 1:**
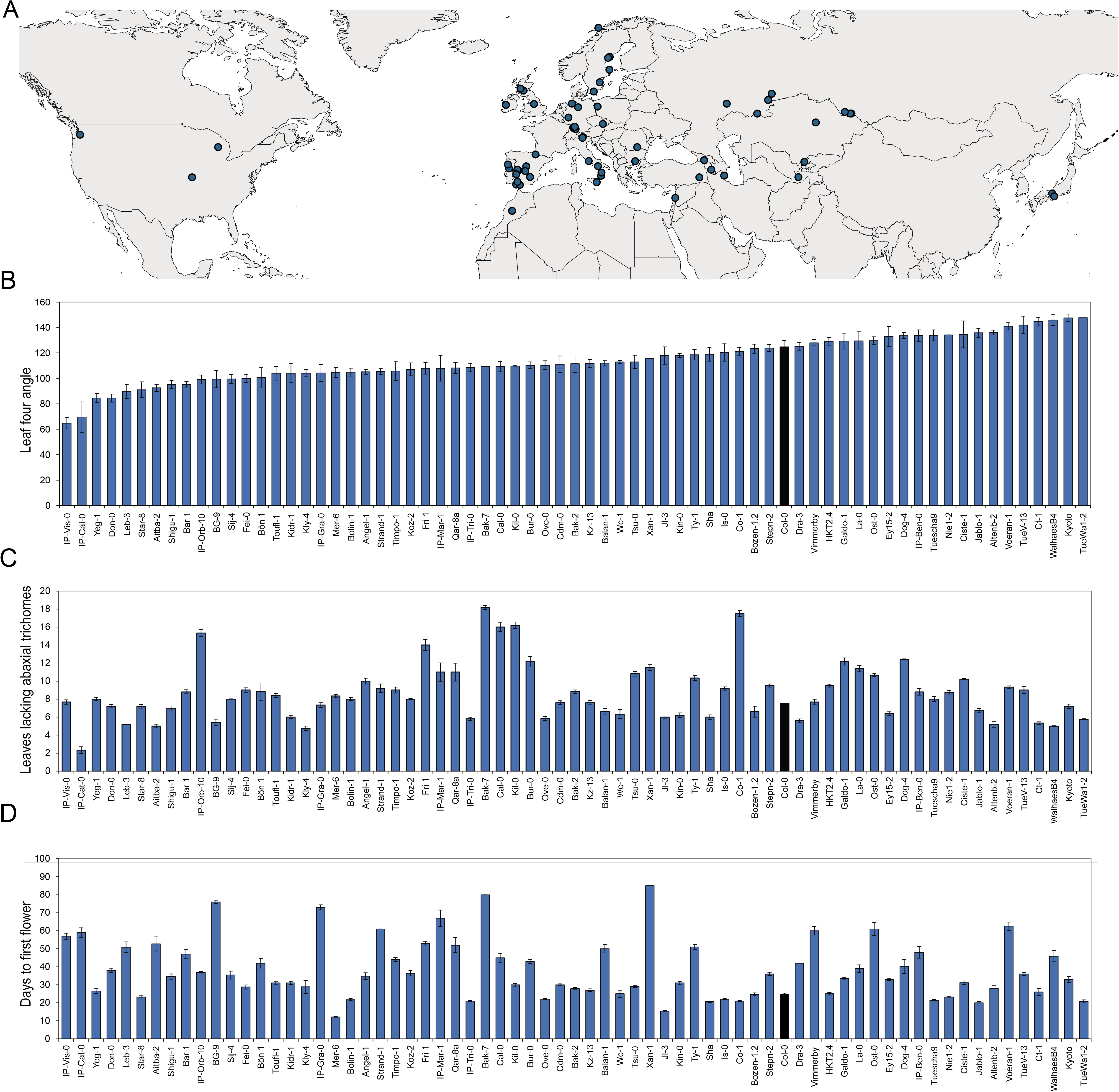
Phenotypic analysis of *A. thaliana* natural accessions. (A) Collection locations of the 70 natural accessions used in this study. (B) Leaf four base angle of natural accessions grown in SD conditions. (C) Number of juvenile leaves, indicated by leaves lacking abaxial trichomes, in natural accessions grown SD conditions, in the same order as (B). (D) Days to first flower of natural accessions grown in LD, in the same order as (B).

We found that accessions varied widely in their leaf four base angle in both SD (Fig 1B) and LD conditions (**Fig. S1A)**, and that the appearance of abaxial trichomes varied from leaf 3 to leaf 18. Although the angle of the leaf base was strongly correlated in SD and LD (**Fig. S1B**), there was no correlation between the angle of the leaf base and the first leaf with abaxial trichomes (Fig. 1B-C**, Fig. S1B-C**). Additionally, there was no relationship between angle of the leaf base and days to first open flower in either SD or LD (Fig. 1B,D **, Fig. S1D**). This suggests that traits associated with timing of vegetative phase change, leaf angle and abaxial trichomes, are regulated by at least partially different mechanisms in these accessions, and that these traits are regulated independently from timing of the reproductive phase.

### Accessions from Central Asia produce adult leaves precociously

Within this group of accessions, we focused on a group of nine accessions from Central Asia that had a similar early adult leaf-shape phenotype (Fig. 2A,D). Principal component analysis (PCA) performed on 19 bioclimatic variables plus altitude from the WorldClim database revealed that these Central Asian accessions were collected from regions with similar climates. This region was positively correlated with temperature seasonality and temperature range, two measurements of high variation in temperature, (Fig. 2B-C) and suggests that these environmental factors promote the development of adult leaf shape. However, as we saw in the larger group of accessions, appearance of early adult leaf shape and production abaxial trichomes were de-coupled. Only four of these accessions produced abaxial trichomes significantly earlier than Col-0, whereas the others produced abaxial trichomes at the same time or slightly later than Col-0 (Fig. 2E-F). This provides further evidence that different phase-specific traits are regulated by different mechanisms in *A. thaliana*.

**Figure 2:**
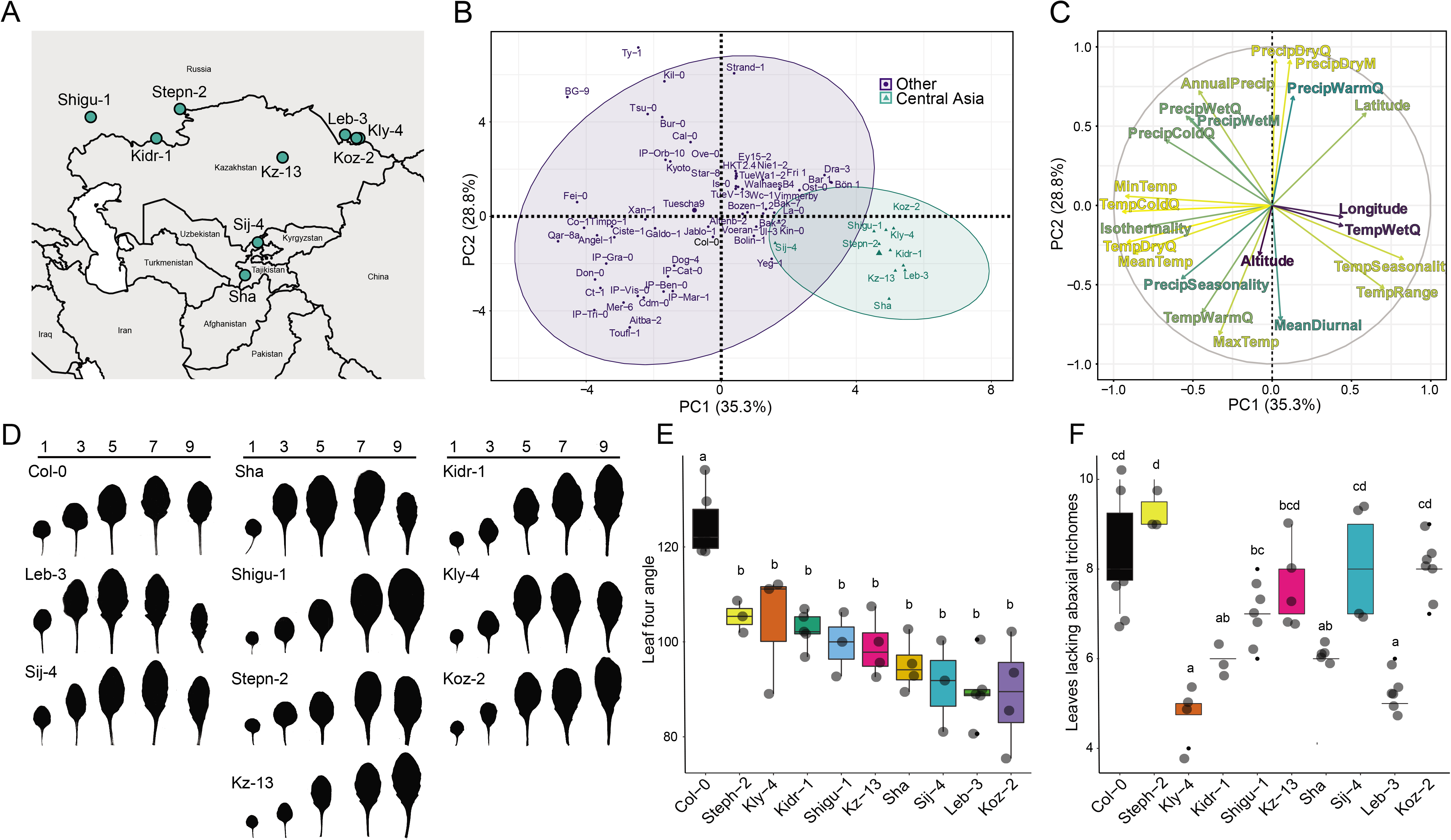
Phenotypic analysis of natural accessions from Central Asia that undergo early vegetative phase change. (A) Collection locations of natural accessions from Central Asia with early adult leaf shape. (B-C) PCA score plot (B) and loading plot (C) for 19 bioclimatic variables and altitude for collection locations of 70 *A. thaliana* accessions. Percentages of variance indicated on each PCA axis. Accessions colored to show Central Asian accessions. (D) Fully expanded rosette leaves 1,3,5,7, and 9 of natural accessions from Central Asia under SD conditions. (E-F) Leaf four base angle (E) and the number of leaves lacking abaxial trichomes (F) of early Central Asian natural accessions. (n=3-6 for each genotype). Significantly distinct groups (denoted by letters) were determined by one-way ANOVA followed by Tukey HSD test (*p*=0.05). Grey dots indicate biological replicates; black dots indicate outliers, center line marks the median value; boxes outline the first and third quartiles; whiskers mark minimum and maximum values.

### Variation in miR156/SPL gene expression does not fully explain natural phenotypic variation in vegetative phase change in Central Asian accessions

To investigate the role of miR156 in these accessions, we measured the abundance of the mature miR156 transcript and the primary transcripts of *MIR156A* and *MIR156C*— which account for about 80% of miR156 pool in in Col-0—in the primordia of leaves 1&2. We found that the level of the mature miR156 was decreased in about half of the Central Asian accessions that showed early leaf shape development, specifically, Leb-3, Stepn-2, Sha and Kly-4. However, the other five accessions had Col-0-like or slightly higher abundance of miR156, which is the opposite of the expected result as high levels miR156 promote juvenile traits (Fig. 3A). Notably, the accessions that had both early leaf shape and abaxial trichome development—Kly-4, Shigu-1, Kidr-1 and Sha—did not all have decreased levels of mature miR156 relative to Col-0. Furthermore, the expression of pri-*miR156A* and pri-*miR156C* is not correlated with the timing of the vegetative phase transition, or with the level of the mature miR156 transcript (Fig 3B-C). For example, Koz-2 had approximately the same amount of miR156 in leaves 1&2 as Col-0, but both pri-*miR156A* and pri-*miR156C* were significantly lower in this accession than in Col-0. Shigu-2 also had approximately the same amount of miR156 as Col-0, but had less pri-*miR156A* and more pri-*miR156C* than Col-0.

**Figure 3:**
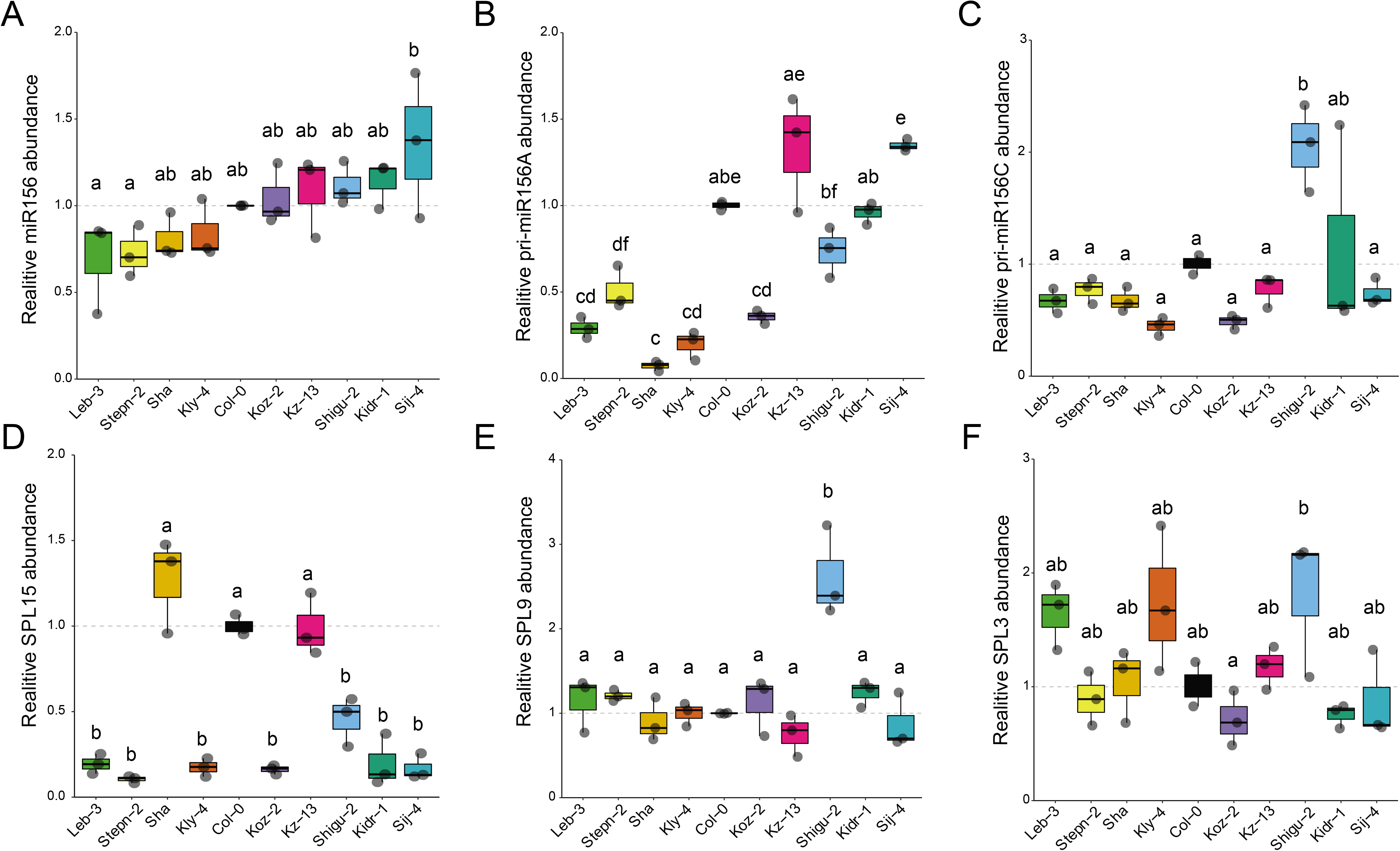
Relative abundance of miR156 and SPL transcripts in leaf primordia of early Central Asian accessions. (A) Mature miR156 abundance relative to SnoR101 in the primordia of leaves 1&2 in Central Asian accessions. (B-F) The abundance of the pri-*miR156A* and pri-*miR156C* primary transcripts, and *SPL15, SPL9, SPL3* mRNAs relative to *ACT2* in the primordia of leaves 1&2 of Central Asian accessions. Accessions are in the same order as in (A). Data are presented as the mean ± SEM (n = 3). Significantly distinct groups (denoted by letters) were determined by one-way ANOVA followed by Tukey HSD test (*p*=0.05). Grey dots indicate biological replicates, black dots indicate outliers, center line marks the median value; boxes outline the first and third quartiles, whiskers mark minimum and maximum values.

To determine if the precocious phase change phenotype of these accessions is regulated downstream of miR156, we measured the abundance three miR156 targets— *SPL15, SPL9, SPL3—*in the primordia of leaves 1&2. There was no clear relationship between the abundance of these transcripts and the expression of miR156 or their phase change phenotype (Fig. 3 D-F). Seven of the nine lines had significantly less *SPL15* levels and eight of the nine lines had *SPL9* was present in approximately the same amount as in Col-0, even though many of these accessions had lower levels of miR156 than Col-0. Shigu-2 had elevated levels of *SPL3* and *SPL9* compared to Col-0, which correlates with its early phase change phenotype, but not with the amount of miR156 in this accession.

These data indicate that regulation of phase-specific aspects of leaf shape and the production of abaxial trichomes is complex, even among accessions with similar geographic and climatic origins and vegetative phase change phenotypes. While some of these accessions—such as Sha and Leb-3—have the expected decreased levels of miR156 or elevated *SPL* gene expression, the early vegetative phenotype of many of these accessions did not reflect the abundance of miR156 or *SPL* transcripts. This suggests that natural variation in phenotypic traits associated with vegetative phase change is controlled by unknown genetic loci in addition to the genes that encode miR156 and *SPL* transcription factors.

### Recombinant inbred lines from Sha x Col-0 reveal eight QTLs that regulate vegetative phase change

Of the accessions from Central Asia, Sha was one of the most consistently precocious in that it had an earlier change in leaf morphology, fewer leaves lacking abaxial trichomes and an earlier increase in leaf serrations compared with Col-0 (Fig. 4A-E). Despite this precocious vegetative phenotype, Sha flowered slightly later than Col-0 after four weeks of vernalization (Fig. 8A,D). Additionally, the early vegetative phase change phenotype of Sha persisted in the absence of vernalization, which significantly delayed its flowering (**FigS2. 2A-C**). This demonstrates that vegetative phase change and floral induction are regulated independently in this accession.

**Figure 4:**
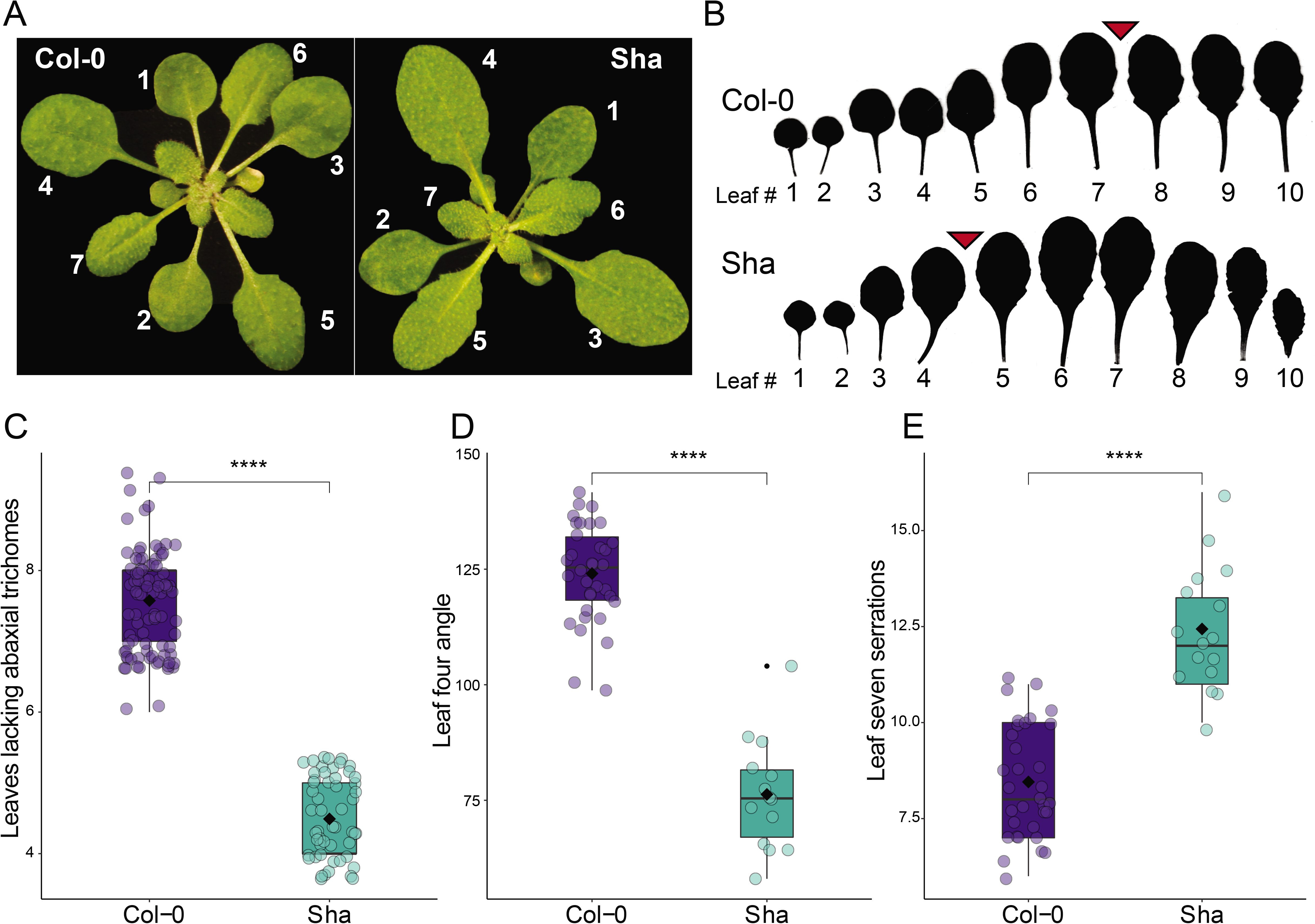
Sha undergoes early vegetative phase change. (A) Four-week-old Col-0 and Sha plants grown in SD. (B) Fully expanded rosette leaves of Sha and Col-0. Red arrows indicate average of the first leaf producing abaxial trichomes. (C-E) The number of leaves lacking abaxial trichomes (C), the angle of the leaf base of leaf four (D), and the number of serrations on the lamina of leaf seven (E) in Sha and Col-0 grown under SD conditions. Significance was determined by Student’s *t-*test (**** = *p*< 0.0001). Blue/green dots indicate biological replicates; black dot indicate outliers; center line marks the median value; boxes outline the first and third quartiles; whiskers mark minimum and maximum values.

To examine the basis for this phenotype, we measured transcript levels of miR156, pri-*miR156A*, pri-*miR156C* in Sha in 2-week-old seedlings, and in the primordia of leaves 1&2 and leaves 5&6 (Fig. 3A). As expected, Col-0 and Sha had relatively high levels of these transcripts in seedlings and leaves 1&2, and had much lower levels in leaves 5&6 (Fig. 5A-C). Sha had significantly less mature miR156, pri-*miR156A*, and pri-*miR156C* than Col-0 in seedlings and leaves 1&2, but had essentially the same amount of these transcripts as Col-0 in leaves 5&6, which are produced after Sha has undergone vegetative phase change. This result suggests that the precocious phenotype of Sha is attributable to an accelerated decline in the expression of *MIR156A* and *MIR156C*, rather than to a decrease in the overall expression of these genes. Additionally, the observation that leaf five has the same level of miR156 in Sha Col-0 yet has a more adult phenotype could also suggest that Sha has a lower sensitivity to miR156. Downstream of miR156, *SPL15* was expressed at a significantly higher level in Sha compared with Col-0 (Fig. 5D). *SPL9* transcripts were consistently, but not significantly, higher in Sha in each of the three samples we examined, but *SPL3* was expressed at similar levels in Sha and Col-0 (Fig. 5E-F). Taken together, these results suggest that the miR156/*SPL* pathway plays a role in the early phase change phenotype of Sha, and made it a promising candidate for further quantitative trait loci (QTL) analysis.

**Figure 5:**
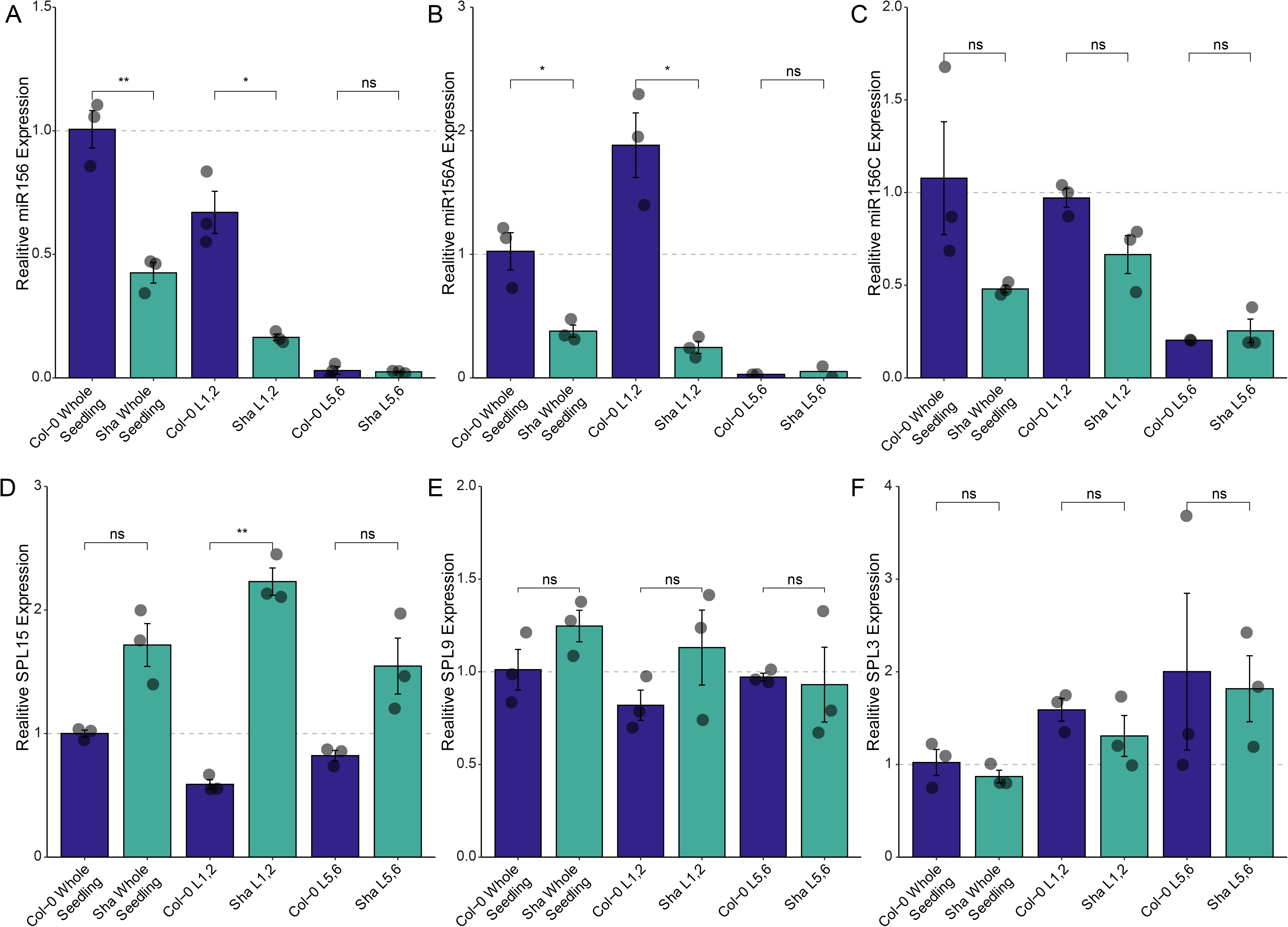
Relative abundance of miR156 and SPL transcripts in Col-0 and Sha. (A) Mature miR156 abundance relative to SnoR101 in seedlings and leaf primordia of Sha and Col-0. Data presented as the mean ± SEM (n = 3). Grey dots indicate biological replicates. Significance was determined by Student’s *t*-test (** = p< 0.01, * = p< 0.05). (B-F) pri-*miR156A*, pri-*miR156C, SPL15, SPL9*, and *SPL3* mRNA abundance relative to *ACT2* in whole seedling and in leaf primordia of Sha and Col-0. Data presented as the mean ± SEM (n = 3). Grey dots indicate biological replicates; black dot indicate outliers. Significance was determined by Student’s *t*-test (** = p< 0.01).

To do this, we took advantage of a set of 164 genotyped recombinant inbred lines (RILs) generated from the F2 progeny of a cross between Sha and Col-0 (Fig. 6A.C) (Simon *et al.* 2008). Ten plants of each line were grown in SD conditions and scored for the leaf three base angle, the number of leaves lacking abaxial trichomes, and the number of serrations on leaf seven. Phenotypic data were analyzed using the R/qtl package (Broman *et al.* 2003).

**Figure 6:**
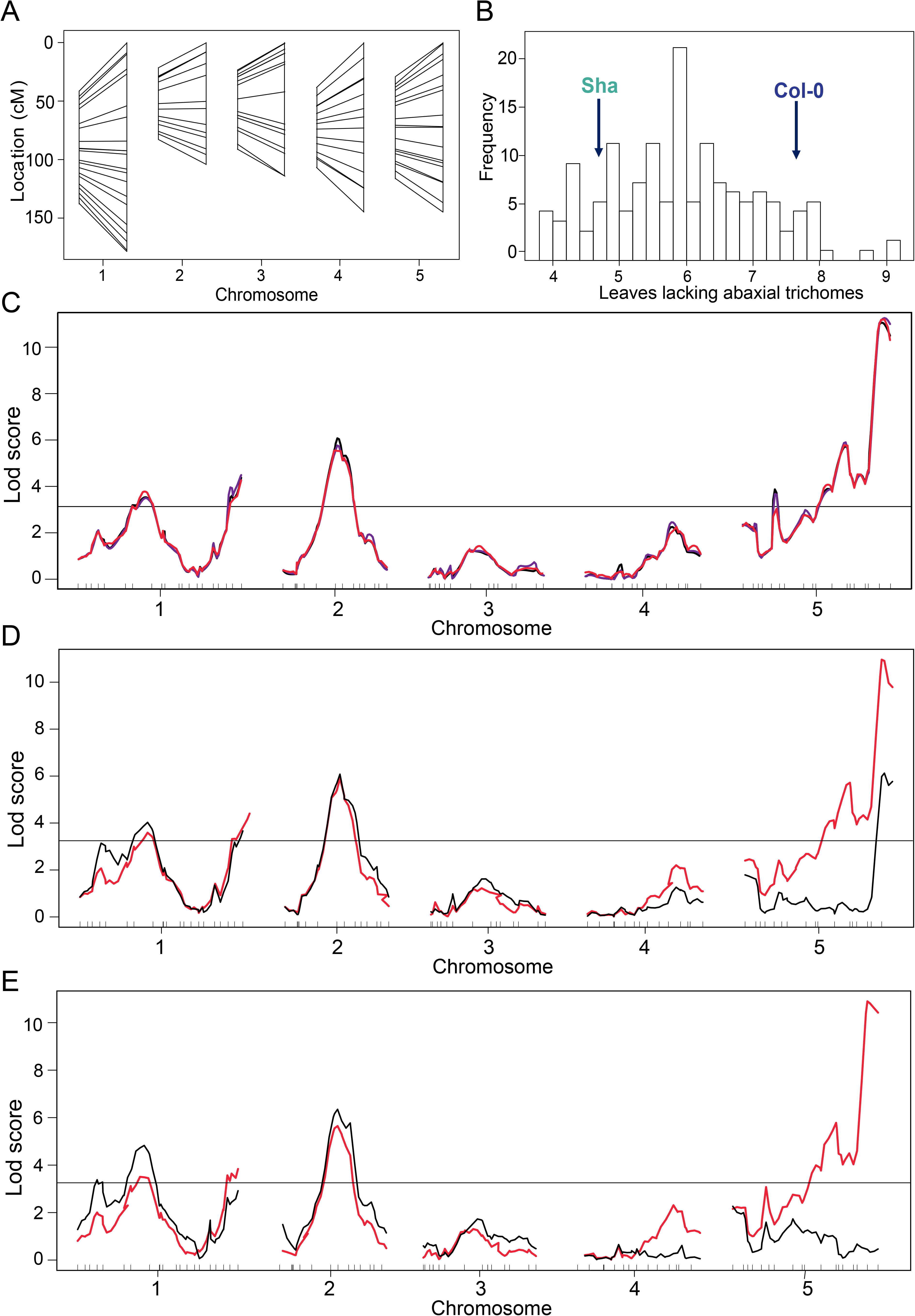
QTL mapping of the number of leaves lacking abaxial trichomes in Sha x Col-0 RILs in SD conditions. (A) Location of molecular markers on the recombination (left) and physical (right) maps in Sha x Col-0 RILs. (B) Phenotypic distribution in the number of leaves lacking axial trichomes of 164 Sha x Col-0 RILs grown in SD conditions. Average phenotype of Col-0 and Sha parents indicated by arrows. (C) QTL analysis for the number of leaves lacking abaxial trichomes with a LOD score threshold of 3.2 based on 1000 permutations and 95% significance. Black lines indicate the LOD Score based on maximum likelihood via the EM algorithm, blue lines indicate Haley-Knott regression, red lines indicate multiple imputation method. (D-E) Covariate analysis using the peak coordinates of the QTLs on chromosome five. Red lines are LOD scores based on leaves lacking abaxial trichomes using maximum likelihood method via the EM algorithm. Black lines are LOD scores based on a genome scan for number of leaves lacking abaxial trichomes with the effects of each QTL peak on chromosome five incorporated as an interacting covariate.

QTL analysis of number of leaves lacking abaxial trichomes, revealed four significant loci using a confidence interval of 95% (Fig. 6C). The most robustly significant QTL was on the bottom of chromosome five, and it accounted for 20.45% of the variance in the whole RIL population. A second significant locus on chromosome two accounted for 10.84% of the variance. Two less significant QTLs were found on chromosome one, accounting for about only ∼3% of the variance (Fig. 6C). Covariance and QTL interaction analysis revealed the QTL on chromosome five was composed of at least two interacting epistatic loci (Fig. 6D-E), but the QTLs on chromosomes one and two were mostly independent of each of the others (**SupFig3A-B**).

QTL analysis of other traits associated with the timing of vegetative phase change produced slightly different results from those found with number of leaves lacking abaxial trichomes. The most significant peak for the leaf three base angle was on chromosome two, which resembled the QTL associated with leaves lacking abaxial trichomes. However, we found a second QTL for this trait on the top of chromosome one, and no QTLs on chromosome five (Fig. 7A-B). An analysis of the number of the number serrations on leaf seven revealed a single significant QTL on the top of chromosome five, in a different location from the QTL for leaves lacking abaxial trichomes (Fig. 7C-D). These results indicate that multiple loci contribute to the precocious vegetative development of Sha, with different loci controlling different aspects of this phenotype.

**Figure 7:**
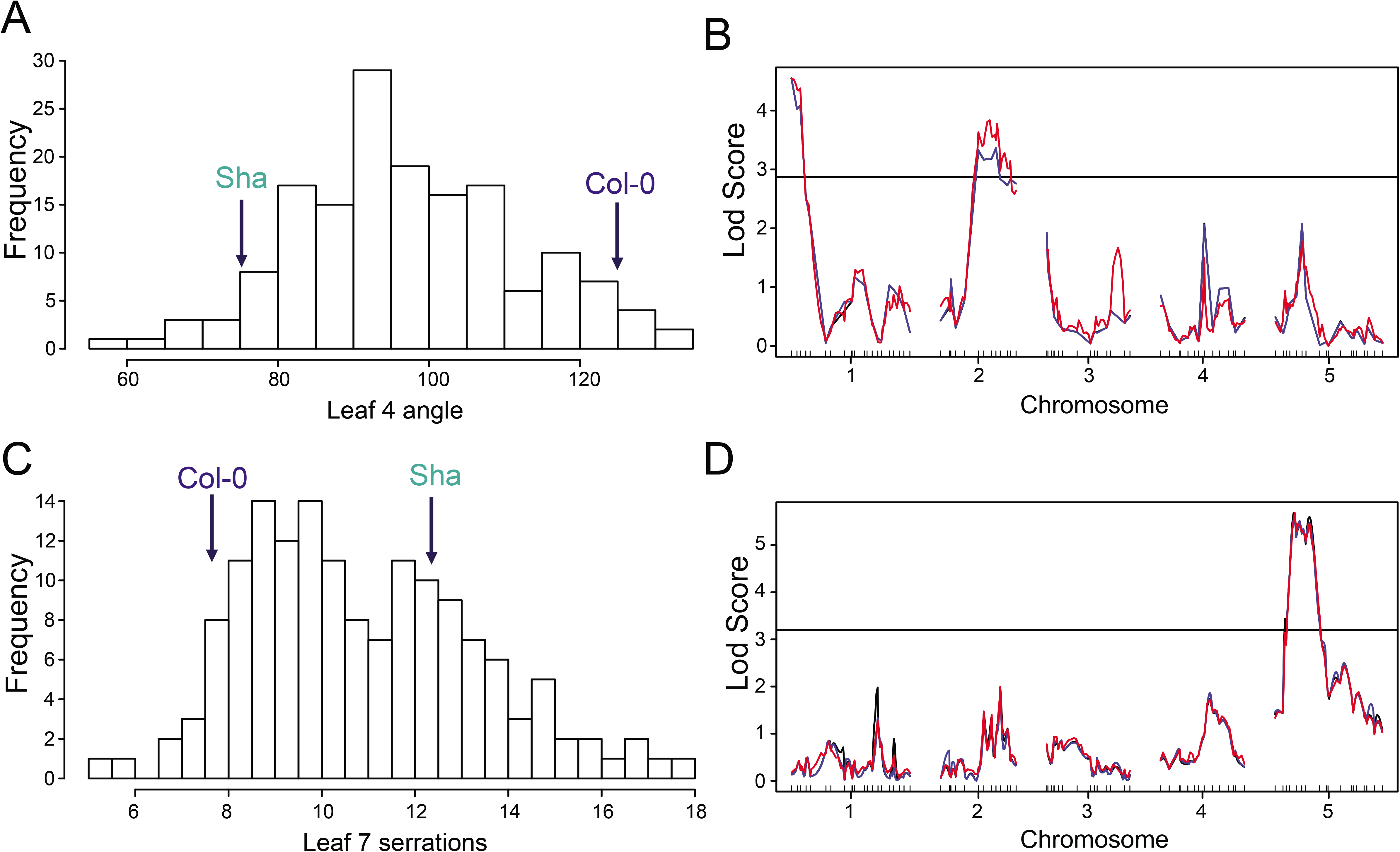
QTL mapping of leaf shape in Sha x Col-0 RILs in SD conditions. (A) Phenotypic distribution in leaf four base angle of 164 Sha x Col-0 RILs grown in SD conditions. Average phenotype of Col-0 and Sha parents indicated by arrows. (B) QTL analysis for leaf four base angle with a LOD score threshold of 2.91 based on 1000 permutations and 95% significance. Black lines indicate the Lod Score based on maximum likelihood via the EM algorithm, blue lines indicate Haley-Knott regression, red lines indicate multiple imputation method. (C) Phenotypic distribution in leaf seven serrations of 164 Sha x Col-0 RILs grown in SD conditions. Average phenotype of Col-0 and Sha parents indicated by arrows. (D) QTL analysis for number of serrations on leaf seven in SD conditions with a LOD score threshold of 3.07 based on 1000 permutations and 95% significance. Black lines are the Lod Score based on maximum likelihood via the EM algorithm, blue lines are Haley-Knott regression, red lines are multiple imputation method.

### RILs grown under long days demonstrate that QTLs regulating vegetative phase change and reproductive phase change are distinct in Sha

To determine if the QTLs that regulate vegetative phase change are independent of the QTLs that regulate flowering time, Sha x Col-0 RILs were also grown under floral inductive (LD) conditions. Ten plants from each genotype were scored for the time to first open flower and for the number of leaves lacking abaxial trichomes. Under LD conditions, Sha produced significantly fewer leaves without abaxial trichomes (Fig. 8A, B), but flowered slightly later (Fig. 8D, E) than Col-0. QTL analysis of the number of leaves lacking abaxial trichomes revealed a QTL at the bottom of chromosome 5, located in the same position as the QTL associated with this trait in SD conditions, as well as less significant QTLs that resembled the QTLs previously detected on chromosome one (Fig. 8C). We did not detect the significant QTL for abaxial trichome production on chromosome two that we had identified under SD conditions (Fig. 8C). QTL analysis of flowering time, using days to first open flower as the phenotype, revealed two QTLs different from those regulating vegetative phase change. The most significant is located on chromosome one, and likely corresponds to the flowering time gene *FT.* Another located at the top of chromosome three that may correspond to the flowering time gene *vrn1* (Fig. 8D-F) (Schwartz *et al.* 2009). These results indicate that flowering time and the timing of vegetative phase change are regulated by different loci in Sha and Col-0. They also demonstrate that the genes that regulate the timing of abaxial trichome production are sensitive to photoperiod and/or light quantity.

**Figure 8:**
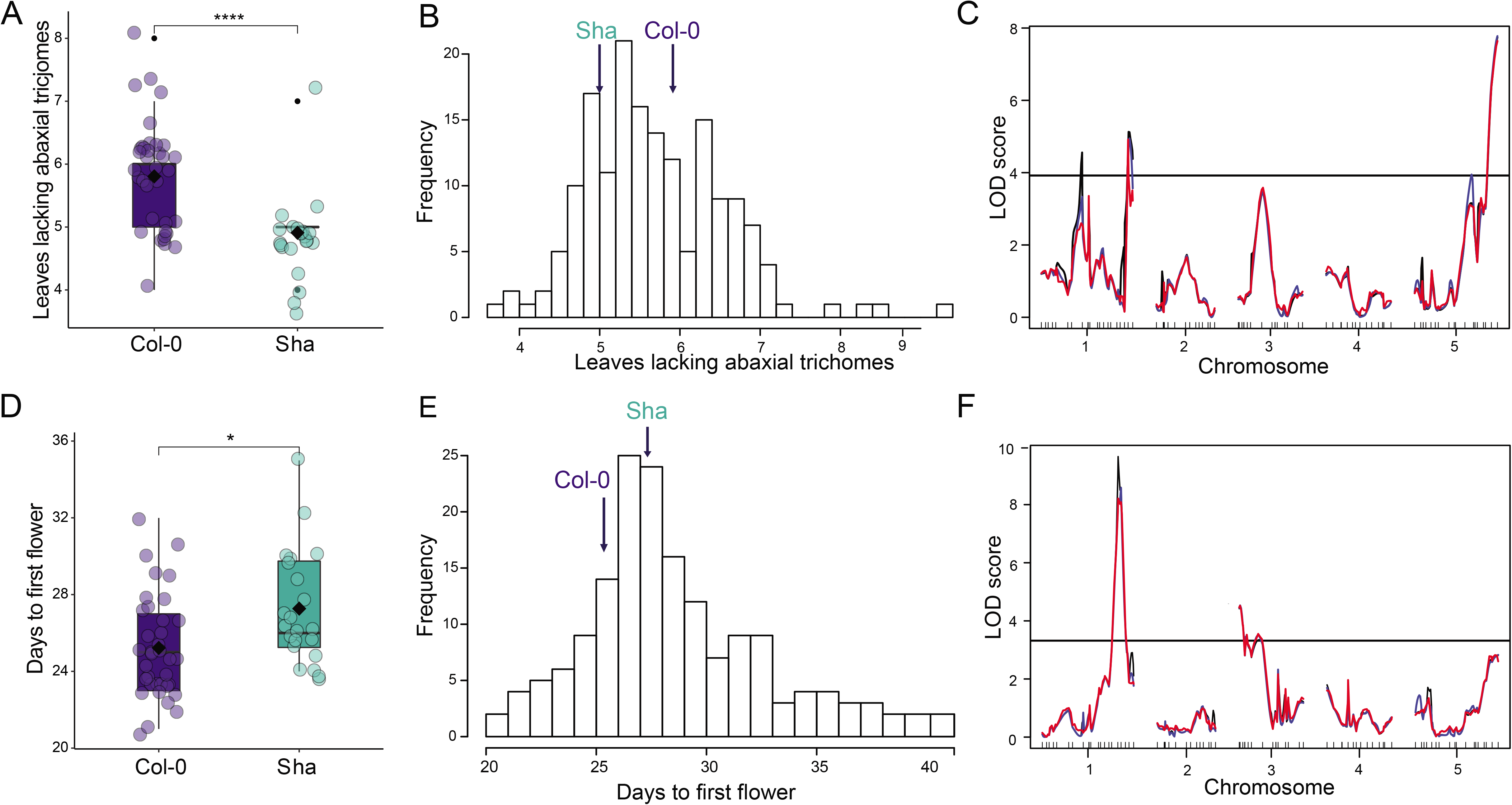
QTL analysis of days to first flower in Sha x Col-0 RILs in LD conditions. (A) The number of leaves lacking abaxial trichomes of Col-0 and Sha grown in LD conditions. Blue/green dots indicate biological replicates; black dot indicate outliers; center line marks the median value; boxes outline the first and third quartiles, whiskers mark minimum and maximum values. (B) Phenotypic distribution in the number of leaves lacking abaxial trichomes of 164 Sha x Col-0 RILs grown in LD conditions. Average phenotype of Col-0 and Sha parents indicated by arrows. (C) QTL analysis for the number of leaves lacking abaxial trichomes grown in LD conditions with a LOD score threshold of 3.92 based on 1000 permutations and 95% significance. Black lines are the Lod Score based on maximum likelihood via the EM algorithm, blue lines are Haley-Knott regression, red lines are multiple imputation method. (D) The days to first open flower in Col-0 and Sha grown in LD conditions. Blue/green dots indicate biological replicates; black dot indicate outliers; center line marks the median value; boxes outline the first and third quartiles; whiskers mark minimum and maximum values. (E) Phenotypic distribution in the number of days to first open flower of 164 Sha x Col-0 RILs grown in LD conditions. Average phenotype of Col-0 and Sha parents indicated by arrows. (F) QTL analysis for days to first open flower grown in LD conditions. LOD score threshold is 3.31 based on 1000 permutations and 95% significance. Black lines are the LOD Score based on maximum likelihood via the EM algorithm, blue lines are Haley-Knott regression, red lines are multiple imputation method.

### Analysis of candidate genes within the Sha x Col-0 QTLs

To identify candidate genes of interest within the QTLs we found, we used available genomic and transcriptomic data from Sha and Col-0 accessions from the 1001 Genomes database (1001genomes.org, 1001epigenomes.org). We focused on the QTLs regulating abaxial trichome development on chromosomes two and five, as they were either identified in analyses in both SD and LD or with other traits associated with vegetative phase change. Both were distinct from the QTLs associated with flowering time.

First, using a *de novo* genomic assembly of Sha (Jiao and Schneeberger 2020), and the Tair10 Col-0 assembly, genomic DNA sequences that contained only the QTLs on chromosome two and chromosome five were used to determine variation in SNPs between Sha and Col-0. From this, the program SNPeff (Cingolani *et al.* 2012) was used to find predicted impact of these SNPs on genes, which totaled over 5,000 protein coding genes (**results shown in Table S1**). In addition, genes within these QTLs were analyzed using RNA-seq data from 1001Genomes database (Kawakatsu *et al.* 2016) to find genes differentially expression between Sha and Col-0 (**results shown in Table S1**). While these analyses are not fully encompassing for finding casual loci, they helped inform our assessment of candidate genes for the regulation of vegetative phase change in Sha within these QTLs.

In the QTL on chromosome five, the candidate that we explored first was *SPL13*, as *SPL13* is a known target of miR156, and elevated levels of *SPL13* have been shown to regulate abaxial trichome production (Xu *et al.* 2016). However, *SPL13* abundance was lower in Sha when compared with Col-0 in whole seedlings and leaves 5&6 (Fig. 9A), likely because Col-0 contains a recent tandem duplication of *SPL13* that is absent in other accessions, including Sha. Additionally, there were no SNPs in Sha with predicted high impact on *SPL13* function, and only two SNPs with predicted low to moderate impact (**Table S1**). As a result, we ruled out *SPL13* as a candidate for regulation of phase change in Sha.

**Figure 9:**
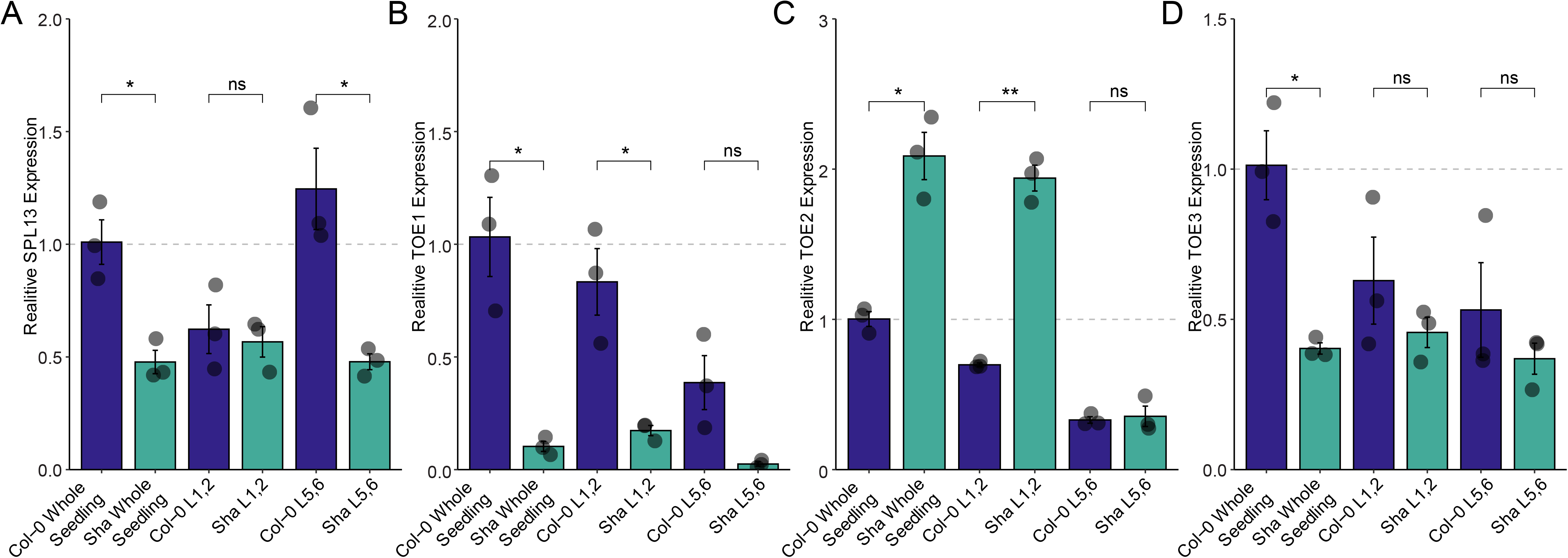
Analysis of candidate genes within significant QTLs on chromosomes two and five. (A-D) RT-qPCR analysis of *SPL13, TOE1, TOE2*, and *TOE3* mRNA abundance relative to *ACT2* in whole seedlings and leaf primordia of Sha and Col-0. Data presented as means ± SEM (n = 3). Grey dots indicate biological replicates, black dot indicate outliers. Significance was determined by Student’s *t*-test (** = p< 0.01, * = p< 0.05).

Two additional candidates within the chromosome five QTL known to regulate abaxial trichome production as well as flowering time are *TOE2* and *TOE3* (Wu et al., 2009). These genes are targets of the microRNA miR172, are highly expressed early in development, and decrease as the plant ages (Aukerman and Sakai 2003; Zhu and Helliwell 2011; Xu *et al.* 2019). We found that *TOE2* was elevated in Sha in 2-week-old seedlings and in the primordia of leaves 1&2, and were present at the same level as Col-0 in leaves 5&6 (Fig. 9C, **Table S1**). This is the opposite the expected result if *TOE2* is responsible for the abaxial trichome phenotype of Sha. *TOE2* has four SNPs within its coding region in Sha, but they were predicted to have low impact on gene function (**Table S1**). *TOE3* was expressed slightly, but not significantly, less in Sha when compared to Col-0 (Fig. 9D, **Table S1**). Furthermore, *TOE3* only has one SNP within its coding region with predicted low impact on gene function. This makes *TOE3* and *TOE2* potential, but unlikely candidates for regulation vegetative phase change in Sha and fine mapping of this QTL will be necessary to determine causal loci in this region.

The chromosome two QTL contained two candidates known to be involved in regulation of phase change: *MIR156A* and *TOE1*. We found pri-*miR156A* levels were much lower in Sha than in Col-0 across development, which matches what would be expected for the precocious phenotype of Sha (Fig. 5A). Similarly, *TOE1*, whose expression promotes juvenile leaf traits, is expressed at a much lower level in Sha than in Col-0 (Fig. 9B, **Table S1**); this is consistent with the early vegetative phase change phenotype of Sha. Moreover, three SNPs within the coding region of *TOE1* are predicted to have a high impact on gene function, and an additional seven SNPs are predicted to have a moderate to low impact on gene function (**Table S1**). Together these data make both *TOE1* and *MIR156A* promising candidates for the QTL on chromosome two.

## Discussion

Juvenile and adult vegetative traits impact plant fitness, and the genetic module that mediates the transition between these phases—miR156 and their *SPL* targets—is conserved across land plants (Axtell and Bowman 2008; Cui *et al.* 2014; Wang and Wang 2015; Leichty and Poethig 2019; Lawrence *et al.* 2021). These observations indicate that vegetative phase change is an important developmental transition in plants, and suggest that an understanding of its genetic architecture will provide insights into both the adaptive significance and molecular mechanism of this transition. Several studies have characterized phenotypic variation in timing of vegetative phase change in natural populations of plants, but few have determined the genetic basis or the adaptative significance of this variation (Wiltshire *et al.* 1998; Jordan *et al.* 2000; Foerster *et al.* 2015). To begin to address these questions, we characterized the timing of vegetative phase change in natural accessions of *A. thaliana*, examined the relationships between leaf development, reproductive development, and climate, and mapped quantitative trait loci affecting phase-specific traits.

Although vegetative phase change is often thought to be linked to reproductive competence (Wang, Czech, and Weigel 2009; Wang 2014; Xie et al. 2020), we observed no correlation between adult vegetative traits (leaf angle or the appearance abaxial trichomes) and reproductive traits (the appearance of the first open flower). We found evidence for this from a comparison of these traits in a worldwide sample of natural accessions and from an analysis of RILs derived from a cross between Col-0 and Sha. Lack of correlation suggests that these developmental transitions are regulated independently and indicates they likely respond differently to natural selection. Furthermore, this emphasizes the importance of taking the timing of vegetative phase change into account when studying natural variation, even in a short-lived plant like *Arabidopsis thaliana*.

Unexpectedly, we found that traits associated with transition to the adult phase, such as appearance of abaxial trichomes and a narrow, serrated leaf shape, were decoupled in many of the natural accessions we studied, including half of the precocious Central Asian accessions. The possibility that these traits are subject to different selective pressures could explain this finding. As trichomes are typically associated with defense against herbivory (Duffey 1986; Dalin *et al.* 2008) and leaf shape is associated with adaptation to temperature and water availability (Givnish 1979; Greenwood 2005), dissociation of these traits may be promoted by varying environmental challenges. This hypothesis supports our finding that seasonal and annual variation in temperature in Central Asia is associated with early development of adult leaf shape, but not production of abaxial trichomes. It also agrees with literature in other species, such as *Eucalyptus* and *Acacia*, where dry and wet climates are associated with variation in the timing of vegetative phase change (Jordan *et al.* 2000; Rose *et al.* 2019).

Consistent with the view that vegetative phase change is regulated by a variety of mechanisms, we found that miR156 and *SPL* expression were not necessarily correlated with the phenotypic variation we observed. This indicates that other mechanisms regulate natural variation in vegetative phase change, in addition to miR156-mediated post-transcriptional repression of *SPL* genes. A variety of factors impact transcriptional regulation of SPLs independent of miR156, including low light intensity (M. Xu, Hu, Poethig 2021) photoperiod (Jung *et al.* 2012) and epigenetic repression by the POLYCOMB REPRESSIVE COMPLEX1 (Picó *et al.* 2015; Li *et al.* 2017). Additionally, the microRNA miR172 regulates abaxial trichome production via its effect on *TOE* gene expression (Wu *et al.* 2009; Huijser and Schmid 2011; Xu *et al.* 2019; Wang *et al.* 2019). Any combination of these mechanisms could be responsible for the variation in vegetative phase change and *SPL* gene expression that we found.

QTL analysis of the basis for the early phase change phenotype of Sha revealed eight QTLs associated with phase-specific traits, only one of which potentially corresponds to genes in the miR156-SPL-miR172-TOE pathway. This provides further evidence that this pathway is not the only source of variation in the timing of vegetative phase change in natural populations. While further work is necessary to find and validate the loci responsible for natural variation in the timing of vegetative phase change, our findings provide the groundwork for such studies. We expect a comprehensive analysis of the genetic basis for natural variation in vegetative phase change will produce new and functionally interesting alleles of genes known to have a role in vegetative phase change, and uncover novel genes involved in the regulation of this developmental transition.

## Data availability

Results from SnpEff and DeSeq of candidate genes are available in **Supplemental Table S1**. All ABRC stock numbers, geographic data and phenotypic data of natural accessions used in this study are available in **Supplemental Table S2**. Sha x Col-0 recombinant inbred lines are available from the Versailles Stock Center at http://publiclines.versailles.inrae.fr/rils/index and genotypic data for these lines is available at http://publiclines.versailles.inrae.fr/page/13. All phenotypic and genotypic data used for QTL mapping are available in **Supplemental Table S3**. All primers used in this study are available in **Supplemental Table S4**.

## Acknowledgments

We thank the Arabidopsis Biological Resource Center and Versailles Stock center for seed stocks, Bishwas Sharma for technical assistance and members of the Poethig lab for helpful discussions.

## Supplemental Material

**Table S1:** SnpEff and DeSeq analysis of candidate genes.

**Table S2:** ABRC catalogue numbers of natural accessions, phenotypic data and climatic data used for this study.

**Table S3:** Phenotypic and genotypic data of Sha x Col-0 RIL lines used for QTL analysis.

**Table S4:** Primers used in this study.

**Figure S1: Leaf angle does not correlate with abaxial trichome production or flowering time.**

(A) Leaf four base angle of natural accessions grown LD conditions, in same order as Figure 1B-D.

(B) Correlation between leaf four angle in LD and SD conditions.

(C) Correlation between leaf four angle and leaves lacking abaxial trichomes in SD conditions.

(D) Correlation between leaf four angle in SD and days to opening of first flower in LD conditions.

(E) Correlation between leaf four angle in SD and days to opening of first flower in LD conditions. Blue lines shown are linear regression model; R-squared and *P*-values from linear regression analysis displayed in upper right corner of each panel.

**Figure S2: Timing of vegetative phase change of Sha and Col-0 in non-vernalized conditions.**

(A-D) The number of leaves lacking abaxial trichomes(A), leaf four angle (B), and days to opening of first flower (C) in non-vernalized SD conditions. Significance was determined by Student’s *t-*test (\*\*\**p*<0.001,\*\*\*\**p*<0.0001).

Blue/green dots indicate biological replicates; black dot indicate outliers; center line marks the median value; boxes outline the first and third quartiles, whiskers mark minimum and maximum values.

**Figure S3: Covariate analysis of leaves lacking abaxial trichomes and QTLs in Sha X Col-0 RILs.**

(A) Covariate analysis using the peak coordinate of the QTL on chromosome two. Red lines are LOD scores based on leaves lacking abaxial trichomes using maximum likelihood via the EM algorithm. Blue lines are the LOD score based on a genome scan for number of leaves lacking abaxial trichomes with the effects of each QTL peak on chromosome five incorporated as an interacting covariate.

(B-C) Covariate analysis using the peak coordinates of the QTLs on chromosome one. Red lines are LOD scores based on leaves lacking abaxial trichomes using maximum likelihood via the EM algorithm. Blue lines are the LOD score based on a genome scan for number of leaves lacking abaxial trichomes with the effects of each QTL peak on chromosome five incorporated as an interacting covariate.

