## Supplementary material for "QTL analysis of vegetative phase change in natural accessions of *Arabidopsis thaliana*": TableS4.docx

| Table S4: Primers used in this study | | |
| --- | --- | --- |
| smRNA RT-qPCR | | |
| Name | Use | Sequence (5’-3’) |
| miR156 RT | miR156 reverse transcription | GTCGTATCCAGTGCAGGGTCCGAGGTATTCGCACTGGATACGACGTGCTC |
| miR172 RT | miR172 reverse transcription | GTCGTATCCAGTGCAGGGTCCGAGGTATTCGCACTGGATACGACATGCAG |
| miR156 Forward | miR156 qPCR | GCGGCGGTGACAGAAGAGAGT |
| miR172 Forward | miR172 qPCR | CGGCGGAGAATCTTGATGATGC |
| Universal Reverse | miRNA qPCR | GTGCAGGGTCCGAGGT |
| SnoR101 F | SnoR101 qPCR | CTTCACAGGTAAGTTCGCTTG |
| SnoR101 R | SnoR101 RT and qPCR | AGCATCAGCAGACCAGTAGTT |

| mRNA RT-qPCR | | |
| --- | --- | --- |
| Name | Use | Sequence (5’-3’) |
| oligo dT | Reverse transcription | TTTTTTTTTTTTTTTTTTTTT |
| ACT2-F | qPCR | GCACCCTGTTCTTCTTACCG |
| ACT2-R | qPCR | AACCCTCGTAGATTGGCACA |
| SPL3-F | qPCR | ATGAGTATGAGAAGAAGCAAAGCG |
| SPL3-R | qPCR | TCCACTACTACTTGTAGCTTTACCT |
| SPL10-F | qPCR | CAGACAAAGGTGTGGGAGAATGCTC |
| SPL10-R | qPCR | TAGGGAAAGTGCCAAATATTGGCG |
| SPL9-F | qPCR | GGAATTTGACCTAGAGAAAAGGAGTT |
| SPL9-R | qPCR | GCATCACCATTTTCGTAAAGCGAAG |
| SPL13-F | qPCR | GGGTTTTCAAGGTAGCAAATTGCT |
| SPL13-R | qPCR | ACCAACAACATAGCTCTGGCTCTG |
| miR156C-F | qPCR | AAAAGCCTCAGATCTAACTCCAACAC |
| miR156C-R | qPCR | GCGTTTCTCTTAAAATTTGTCCCAAAACT |
| miR156A-F | qPCR | CTTCGTTCTCTATGTCTCAATCTCTC |
| miR156A-F | qPCR | TGATTAAAGGCTAAAGGTCTCCTC |
| TOE1-F | qPCR | CGAGTTATAATAATCCCGCCGAG |
| TOE1-R | qPCR | TTAAGGGTGTGGATAAAAGT |
| TOE2-F | qPCR | ATGGAGAACCACATGGCTGC |
| TOE2-R | qPCR | GGTGCTGTAGCTGCTACGGC |
| TOE3-F | qPCR | CTCACCCGATCATCACGAAG |
| TOE3-R | qPCR | TATTGAAACCGGACCAACGA |
