## Supplementary figures and images for "QTL analysis of vegetative phase change in natural accessions of *Arabidopsis thaliana*"

### FigureS1.pdf

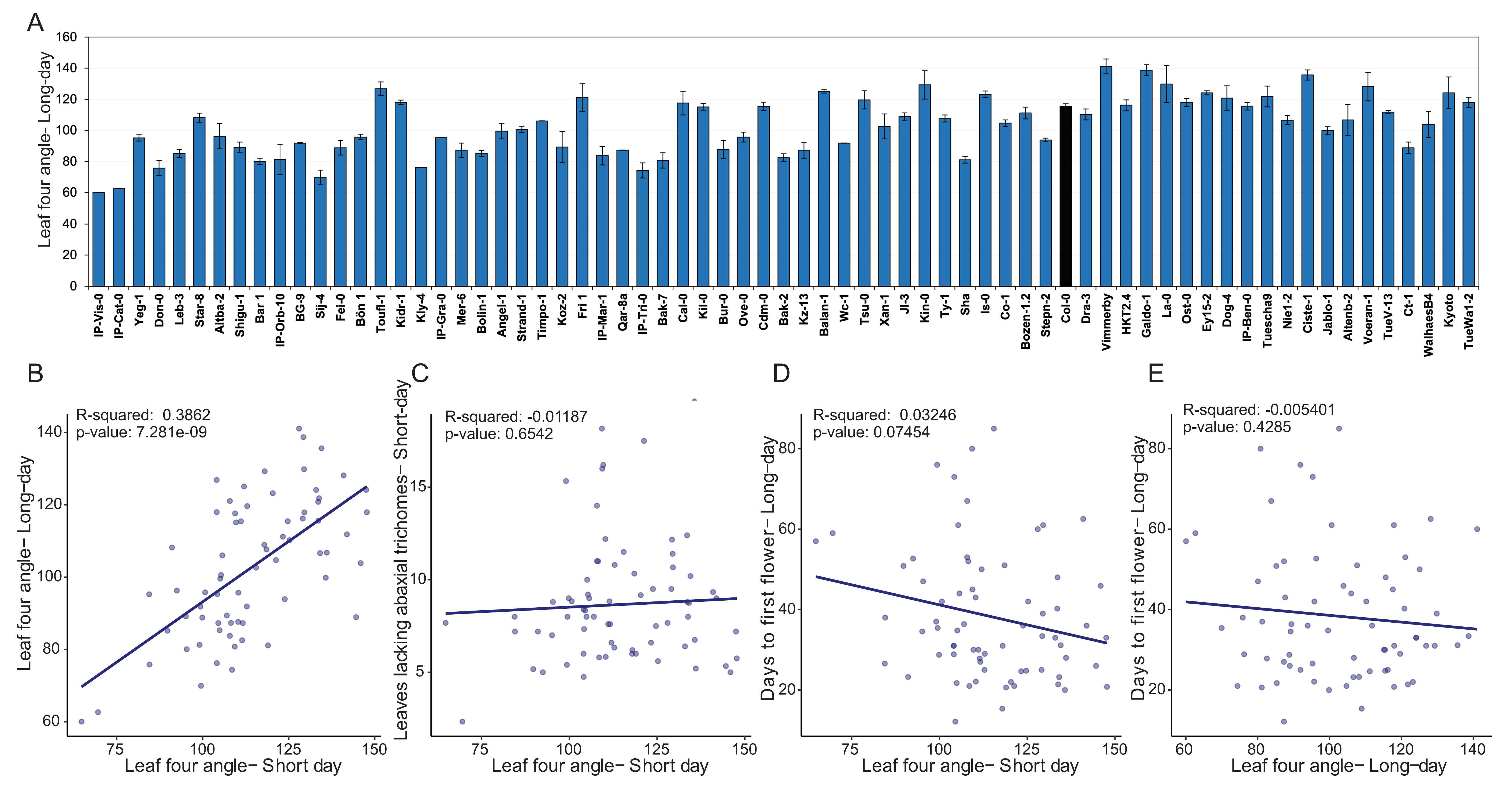

### FigureS2.pdf

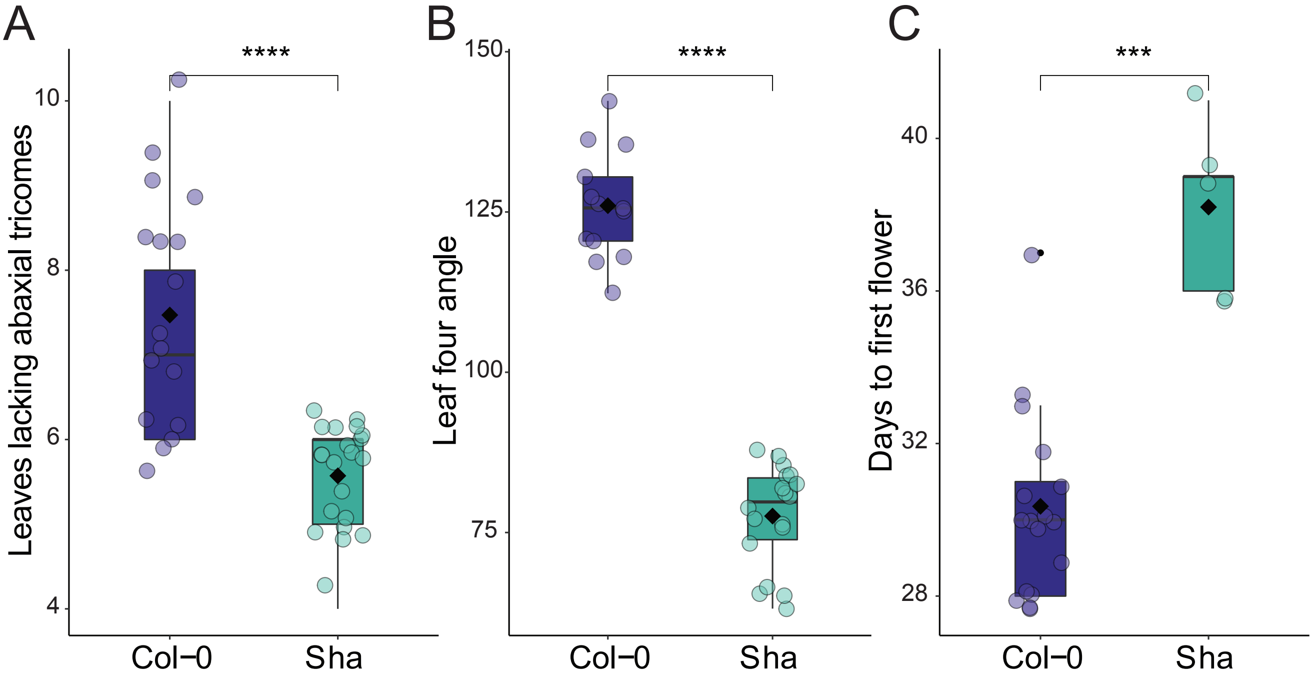

### FigureS3.pdf

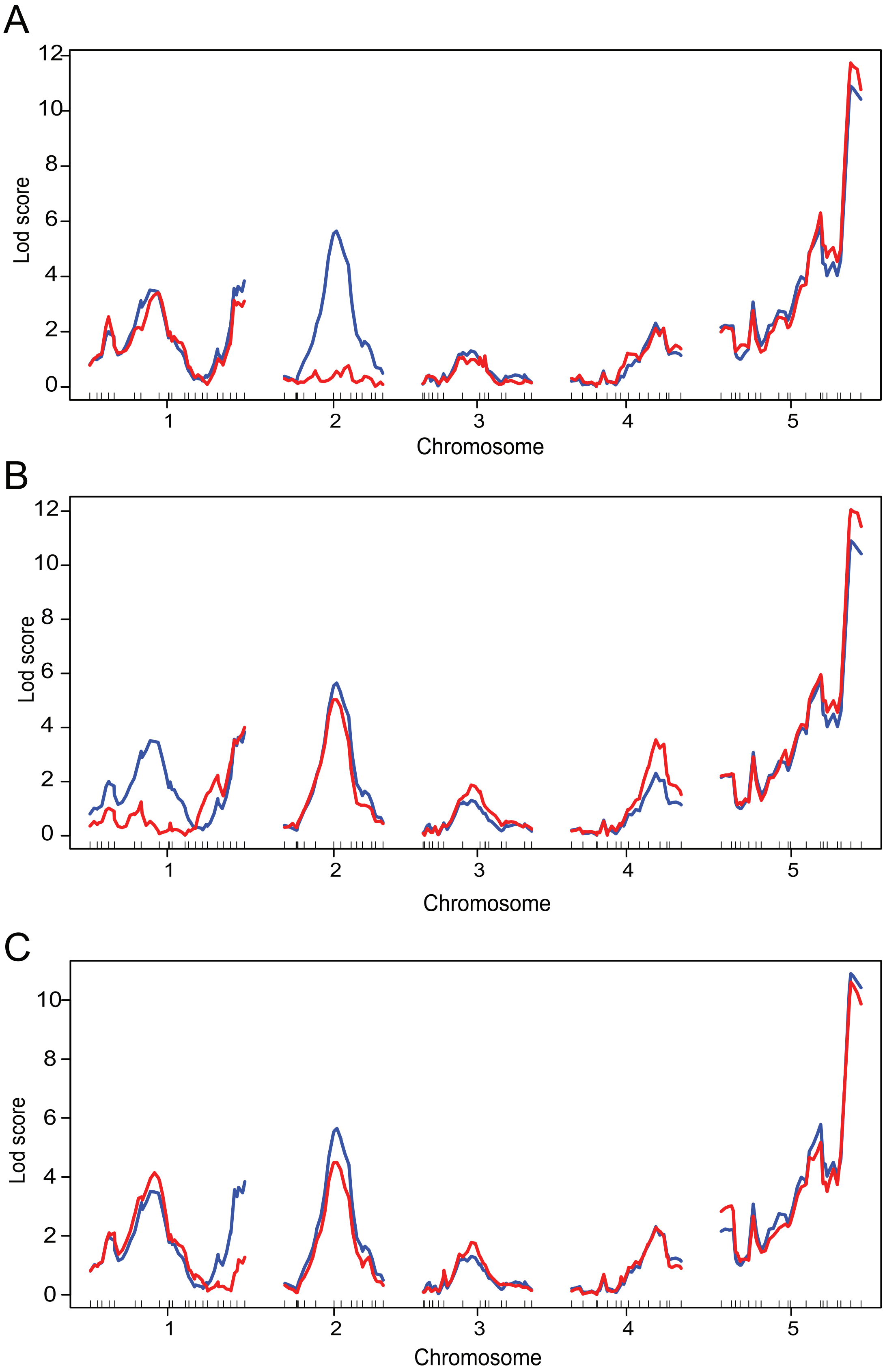
